# Intestinal Transgene Delivery with Native *E. coli* Chassis Allows Persistent Physiological Changes

**DOI:** 10.1101/2021.11.11.468006

**Authors:** Baylee J. Russell, Steven D. Brown, Anand R. Saran, Irene Mai, Amulya Lingaraju, Nicole Siguenza, Erica Maissy, Ana C. Dantas Machado, Antonio F. M. Pinto, Yukiko Miyamoto, R. Alexander Richter, Samuel B. Ho, Lars Eckmann, Jeff Hasty, Alan Saghatelian, Rob Knight, Amir Zarrinpar

## Abstract

Live bacterial therapeutics (LBT) could reverse disease by engrafting in the gut and providing persistent beneficial functions in the host. However, attempts to functionally manipulate the gut microbiome of conventionally-raised (CR) hosts have been unsuccessful, because engineered microbial organisms (i.e., chassis) cannot colonize the hostile luminal environment. In this proof-of-concept study, we use native bacteria as chassis for transgene delivery to impact CR host physiology. Native *Escherichia coli* isolated from stool cultures of CR mice were modified to express functional bacterial (bile salt hydrolase) and eukaryotic (Interleukin-10) genes. Reintroduction of these strains induces perpetual engraftment in the intestine. In addition, engineered native *E. coli* can induce functional changes that affect host physiology and reverse pathology in CR hosts months after administration. Thus, using native bacteria as chassis to “knock-in” specific functions allows mechanistic studies of specific microbial activities in the microbiome of CR hosts, and enables LBT with curative intent.

---

Gut microbes sense and condition the luminal environment over time periods measured in years. Numerous chronic human diseases, including obesity,^1, 2^ non-alcoholic fatty liver disease,^3, 4^ type 2 diabetes,^2, 5^ atherosclerosis,^6-8^ polycystic ovary syndrome,^9^ inflammatory bowel disease,^10^ and cancer,^11^ have protracted symptomology and have consequently been targets of potential engineered, live bacterial therapeutics (LBTs) owing to their long-term capabilities. LBT can also be designed to express functions that address monogenic inborn errors of metabolism (e.g., phenylalanine metabolism in patients with phenylketonuria) and other orphan diseases.^12^ Engineered bacteria also enable otherwise challenging mechanistic studies, allowing researchers to learn how individual bacterial functions or proteins in the microbiome contribute to host physiology, pathophysiology, or the microbiome community.^13^ Although synthetic biologists have designed increasingly elaborate genetic circuits that in principle could risk-stratify patients and combat disease, these circuits have only worked in reduced community models *in vivo* (i.e., gnotobiotic mice, antibiotic treated mice).^14, 15^

A key step in engineered LBT is selection of a microbial host organism, or chassis, which would enable environment-sensing, regulated gene expression, and production of a therapeutic product. Current LBT chassis (e.g., *Escherichia coli* Nissle 1917, *Bacteroides* spp, *Lactobacillus* spp) cannot engraft or even survive in the gut luminal environment. They require frequent re-administration and are difficult to localize to specific regions such as the proximal gut, thereby resulting in unreliable delivery of function.^12, 14, 16^ The lack of engraftment severely limits the use of LBT for chronic conditions, for curative effect, or to help devise tools to study specific functions in the gut microbiome.^14, 16^ The only published human trial using an engineered LBT, where an interleukin-10 (IL-10) expressing *Lactococcus lactis* was used to treat inflammatory bowel disease, yielded disappointing results.^17^ The authors attributed therapeutic failure to the inability of the chassis to survive long enough to express the transgene of interest. Hence, appropriate chassis selection remains a significant barrier to translating synthetic biology applications to human diseases.

LBT face many challenges to survival in the luminal environment, both from the host (e.g., peristalsis, innate and adaptive immunity) and other native microorganisms (e.g., competition, niche availability).^18^ Probiotic strains not adapted to the luminal environment cannot easily compete with microbes that have adapted to their specific luminal niche over thousands of generations.^19^ This phenomenon is apparent in most patients who have received fecal microbiota transplant, where the individual’s native microorganisms return and largely or completely displace the transplanted microbes.^20^ Early studies of native *E. coli* strains, however, demonstrate prolonged colonization and delivery of an inflammation detection circuit.^21^ However, whether native species, particularly native *E. coli*, can be used to functionally change the luminal environment, induce physiological change, or treat disease remains unclear.

In theory, native bacteria are already maximally adapted to the luminal environment of the host, thereby bypassing nearly all the barriers to engraftment, making them an ideal chassis for transgene delivery. The reluctance to use undomesticated native bacteria is driven by the assumption that they are difficult to culture and modify, although recent studies demonstrate that they can be modified more consistently with novel methods.^22, 23^ Studies in gnotobiotic mice show that a host mono-colonized with a single species is resistant to colonization by the same, but not different, species,^24^ suggesting that if the niche of the engineered bacteria is already filled, engraftment would be more difficult. In addition, many have assumed that *E. coli* are not good colonizers due to disappointment with their commensal lab strain counterparts (e.g., *E. coli* Nissle 1917). Lab strain *E. coli* lose colonization or never engraft and lose transgenes of interest or not express them in a way that can affect host physiology. This type of inconsistency makes their use difficult. Their lack of effectiveness has led researchers to favor species that are far more abundant in the gut microbiome (e.g., *Bacteroides* spp, *Lactobacillus* spp.) that are much more challenging to engineer. Hence, it may seem that the barriers to use native bacteria as chassis for transgene delivery, particularly for conventionally-raised (CR) hosts, are insurmountable.

Here we present a proof-of-concept study that advances our ability to use engineered bacteria to effectively change physiology in CR, wildtype (WT) hosts, by demonstrating that native bacteria can serve as chassis for transgene delivery that modifies host phenotype. This is accomplished by identifying a genetically tractable, native bacterial strain from a host (i.e., an undomesticated *E. coli*), modifying this strain to express a transgene of interest, and then reintroducing the engineered native bacteria to the host (**Fig. 1A**). The transgenes of interest included bile salt hydrolase (BSH), a prokaryotic bile acid deconjugation enzyme (**Fig. S1A**) which can potentially affect host metabolic homeostasis, and IL-10, a mammalian cytokine which can function as an anti-inflammatory agent. After a single treatment, engineered native bacteria engraft throughout the entire gut of CR hosts for the lifetime of the mouse, retain the function of their transgene, induce metabolomic changes, affect host physiology, and even ameliorate pathophysiological conditions. We evaluate the robustness of the approach by performing experiments in non-sterile facilities, testing biocontainment between cohoused littermates, and changing diets. We demonstrate the translational potential of this method by successfully and stably transforming native, undomesticated, human-derived *E. coli* that can be used for transgene delivery in humans to potentially treat disease. Although the individual steps of the proposed approach for engineering undomesticated native bacteria to express the desired function of interest are not particularly difficult, the combination is novel, and, put together, they clearly demonstrate that even resource-limited labs can accomplish what has yet to be achieved with other synthetic biology approaches: persistent functional manipulation of the luminal environment of CR hosts to study their physiological effects.

**Figure 1.**
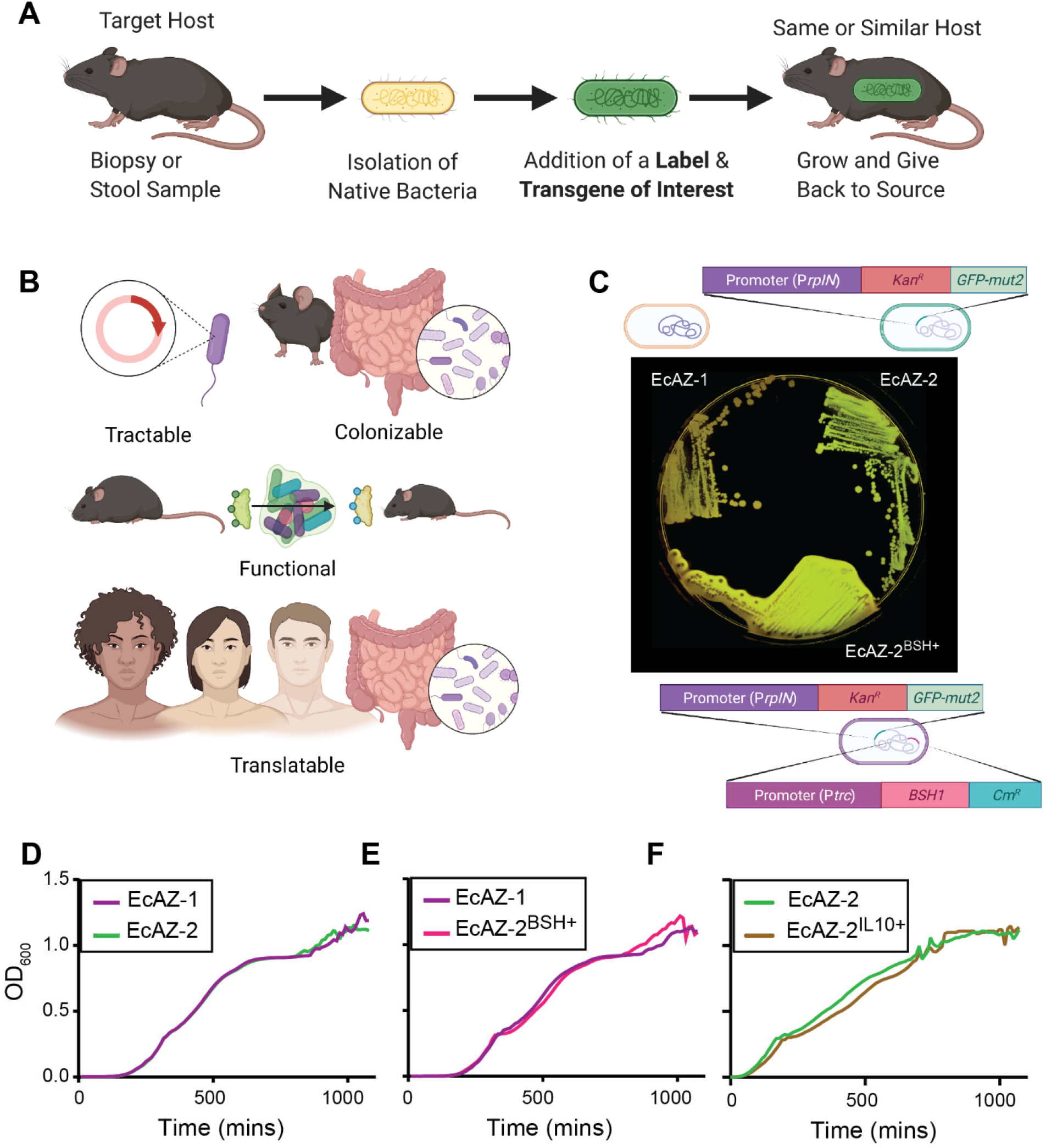
Gut native *E. coli* are genetically tractable and can serve as a chassis for transgene delivery. (**A**) Experimental strategy of engineered native bacteria. (**B**) Optimal characteristics of a chassis for transgene delivery for a potential LBT. (**C**) Original isolate (EcAZ-1), GFP producing strain (EcAZ-2), and GFP and BSH producing strain (EcAZ-2^BSH+^) plated on LB containing lactose and TDCA. Precipitate around EcAZ-2^BSH+^ is the result of TDCA deconjugated by the bacteria to DCA and indicates enzyme functionality. (**D**) Growth curve of EcAZ-2 compared to EcAZ-1. (**E**) Growth curve of EcAZ-2^BSH+^ compared to EcAZ-1. (**F**) Growth curve of EcAZ-2^IL10+^ compared to EcAZ-2. For (D), (E), and (F), the line represents an average of three measurements per strain.

## RESULTS

### Gut native E. coli are genetically tractable

An ideal chassis is genetically tractable, can stably and persistently colonize the selected host, express the gene of interest persistently (and without much effect on bacterial fitness), and impose a functional change to alter the lumen and/or serum to affect host physiology (**Fig. 1B**). Importantly, the strategy should be easily translated to humans since this has been a particular challenge in adapting synthetic biology tools to clinical problems (**Fig. 1B**). In our studies, we used an undomesticated, native *E. coli*, EcAZ-1, isolated from the stool of a CR-WT C57BL/6 male mouse acquired from Jackson Laboratory (Bar Harbor, ME). Green fluorescent protein (GFP) linked with kanamycin resistance was introduced to EcAZ-1 via phage transduction to create a traceable version of the bacteria, EcAZ-2 (**Fig. 1C**). EcAZ-2 was then engineered separately, to form two different engineered native bacteria. The first, EcAZ-2^BSH+^, expresses bile salt hydrolase (BSH; **Fig. S1**), a bacterial gene that causes specific bile acid biotransformations demonstrated to affect the host metabolic phenotype in reduced community microbiome studies (e.g., gnotobiotic mice, antibiotic-treated mice).^25-27^ *In vitro* BSH activity was monitored with the aid of taurodeoxycholic acid (TDCA) plates; as EcAZ-2^BSH+^ deconjugates TDCA to deoxycholic acid (DCA), these plates form a white precipitate around the bacterial colony (**Fig 1C**). The second gene, IL-10 (to form EcAZ-2^IL10+^), is a mammalian anti-inflammatory cytokine. Although continuous administration of IL-10 expressing probiotics has been successful in treating preclinical models of colitis,^28^ this therapy has not successfully translated to humans due the inability of chassis to survive in the luminal environment.^17^ *In vitro* IL-10 production was monitored with an ELISA on bacterial cell lysates. The addition of *gfp* and kanamycin resistance, as well as *bsh* and *Il-10* did not affect the *in vitro* growth rate of EcAZ-2, EcAZ-2^BSH+^, and EcAZ-2^IL10+^ (**Fig. 1D, E, F**). Thus, native *E. coli* are genetically tractable, can express prokaryotic and eukaryotic transgenes, and can potentially serve as chassis for transgene delivery.

### Engineered native E. coli can engraft in the gut lumen

Next, we assessed the candidate chassis’s ability to stably colonize the murine gut. EcAZ-2 stably maintained colonization in 100% (n = 8) of CR-WT hosts, non-antibiotic treated mice for over 110 days in a non-sterile, low barrier mouse facility (**Fig. 2A, S2A**). Domesticated lab bacteria that are often used as chassis for LBT applications (i.e., *E. coli* MG1655 and *E. coli* Nissle 1917) lost colonization over time (p = 0.02, log-rank test, **Fig. S2A**). Mice that maintained colonization had nearly two orders of magnitude lower engraftment (as determined by cfu/g of stool) than those treated with EcAZ-2 (**Fig. 2A**). Colonization of EcAZ-2 was unaffected by changes to the diet macronutrient profile (**Fig. 2B**). Co-housing of mice colonized with EcAZ-2 with non-colonized mice did not lead to horizontal transmission between mice despite coprophagia (**Fig. 2C**). The addition of BSH or IL-10 activity does not significantly alter the colonization level of EcAZ-2; this native strain remains stably colonized in CR-WT mice, for the life of the animal in reproducible experiments (**Fig. 2D, S2B, 2E, S2C**). Sex of the CR-WT host did not affect how well EcAZ-2 colonized the host (**Fig S2D**).

**Figure 2.**
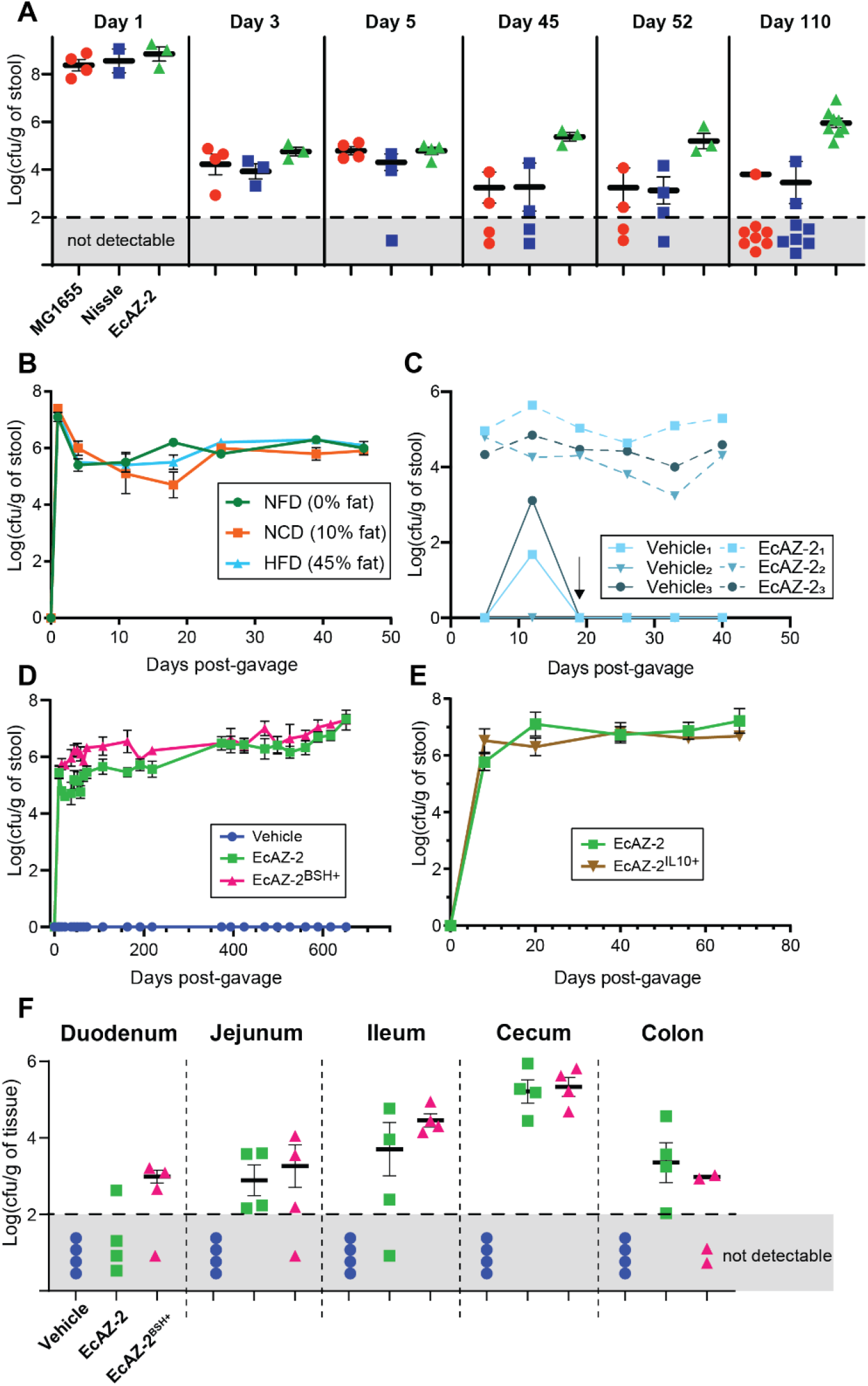
Native E. coli can engraft in the luminal environment. (**A**) Long-term colonization of *E. coli* MG1655 and *E. coli* Nissle 1917 (two lab-strain *E. coli* often used as chassis for function delivery), as well as EcAZ-2 in CR-WT mice housed in a low-barrier, non-sterile facility after a single gavage. Sample measures in the gray area denote mice where the bacteria were not detectable at a limit of detection of Log_10_ 2. (4-8 mice/condition). (**B**) Colonization of EcAZ-2 in stool of CR-WT mice on a stable diet (10% fat), or on diets with different macronutrient profiles (non-fat diet [NFD] or high-fat diet [HFD]). Diet was changed 4 days post-gavage (4 mice/condition). (**C**) Colonization in co-housed mice where one mouse received EcAZ-2 and the other received vehicle via oral gavage. Arrow indicates change in stool collection protocol where mice were separated from their cage mates 24h prior to their weekly stool collection. Mice were returned to their cage mates right after stool collection. Thus, though uncolonized mice are exposed to EcAZ-2 through coprophagia, the level is not sufficient to lead to engraftment. (**D**) Colonization of EcAZ-2 and EcAZ-2^BSH+^ in CR-WT mice housed in a specific-pathogen free facility (4-12 mice/condition). (**E**) Colonization of EcAZ-2 and EcAZ-2^IL10+^ in CR-WT mice housed in a specific-pathogen free facility (4 mice/condition). (**F**) Colonization of gastrointestinal tract in CR-WT mice three months after a single gavage. Sample measures in the gray area denote mice where the bacteria were not detectable at a limit of detection of Log_10_ 2. (4 mice/condition). All error bars indicate standard error of the mean. The marker covers some error bars in panel **B, D**, and **E**.

To determine whether BSH function was retained during engraftment, bacterial isolates from mice treated with EcAZ-2^BSH+^ were grown on TDCA plates. Isolates from every region of the GI tract of every mouse treated with EcAZ-2^BSH+^, across every experiment, retained BSH functionality, confirmed by deconjugation of TDCA to DCA (see example **Fig. S2E**). Likewise, cell lysates of bacterial isolates obtained from mice treated with EcAZ-2^IL10+^ at 89 days after gavage produced IL-10 at far higher levels than the negative control bacteria (i.e., EcAZ-2; **Fig. 2F**).

We cultured gastrointestinal tissues (i.e., duodenum, jejunum, ileum, cecum, and rectum) to determine the extent of bacterial colonization in the various regions of the gut. In contrast to what has been reported for engineered non-native *E. coli*,^12^ EcAZ-2 colonizes the entire GI tract regardless of whether it contains the *bsh* gene, *Il-10* gene, or not, with highest colonization in the cecum and ileum for the BSH strain and cecum and jejunum for the IL-10 strain (**Fig. 2F, S2G, S2H**). Colonization throughout the GI tract was consistent in fed and fasted mice (**Fig. S2G**). These experiments demonstrate that EcAZ-2 can stably and persistently colonize CR-WT hosts for perpetuity while retaining functionality of a gene of interest.

### Engineered native E. coli can perform a luminal function

After demonstrating that native bacteria are genetically tractable and can persistently colonize the entire gut, we assessed whether the transgene delivered by the EcAZ-2 chassis can functionally manipulate the luminal environment of CR hosts. We used *bsh* as our transgene to evaluate the native bacterial chassis, because its functions are well-described and it has been studied in reduced community/gnotobiotic models.^25-27^ To confirm that BSH-expressing native *E. coli* are functional *in vivo*, we determined the effect of EcAZ-2^BSH+^ on the bile acids of mono-colonized gnotobiotic mice. Whereas mice mono-colonized with EcAZ-2 had only primary conjugated bile acids and no deconjugated bile acids (since they lack bacteria that can deconjugate; see methods), mice mono-colonized with EcAZ-2^BSH+^ had greatly reduced primary conjugated fecal bile acids (**Fig. S3A**) and increased primary deconjugated fecal bile acids (**Fig. S3B**) consistent with increased BSH activity. Quantitative analysis of the conjugated and unconjugated individual primary bile acids showed that EcAZ-2^BSH+^ significantly shifted the log ratio of cholic acid to taurocholic acid (CA:TCA) and β-muricholic acid to tauro-β-muricholic acid (bMCA:TbMCA) toward the deconjugated bile acids in mono-colonized mice (**Fig. S3C**).

To determine whether engineered native bacteria expressing BSH can functionally manipulate the fully intact microbiome of CR mice, we gavaged EcAZ-2 or EcAZ-2^BSH+^ into 11-week-old CR-WT mice. Twelve weeks after the single oral administration, stool samples were collected and analyzed for bile acids. CR mice colonized with EcAZ-2^BSH+^ had significantly higher total fecal bile acid levels compared to CR mice colonized with EcAZ-2 (**Fig. 3A**). In addition, CR mice gavaged with EcAZ-2^BSH+^ had decreased primary fecal bile acids (**Fig. 3B**) and decreased primary conjugated bile acids (**Fig. 3C**) compared to mice treated with EcAZ-2. Surprisingly, primary unconjugated bile acids were also significantly decreased in EcAZ-2^BSH+^ treated mice compared to EcAZ-2 controls (**Fig. 3D**). However, this could be due to increased conversion of deconjugated bile acids to secondary bile acids, which was significantly elevated in EcAZ-2^BSH+^ mice (**Fig. 3E**). In fact, treatment with EcAZ-2^BSH+^ affected nearly every fecal bile acid measured, including TbMCA, a potent farnesoid X receptor (FXR) antagonist, and DCA, a potent G protein-coupled bile acid receptor 1 (TGR5) agonist (**Fig. S3D**).^29^ These fecal bile acid data confirm that BSH is functional *in vivo*. Moreover, they demonstrate that native *E. coli* are valuable chassis that can express a function of interest intraluminally at least 12 weeks after a single gavage in CR-WT mice.

**Figure 3.**
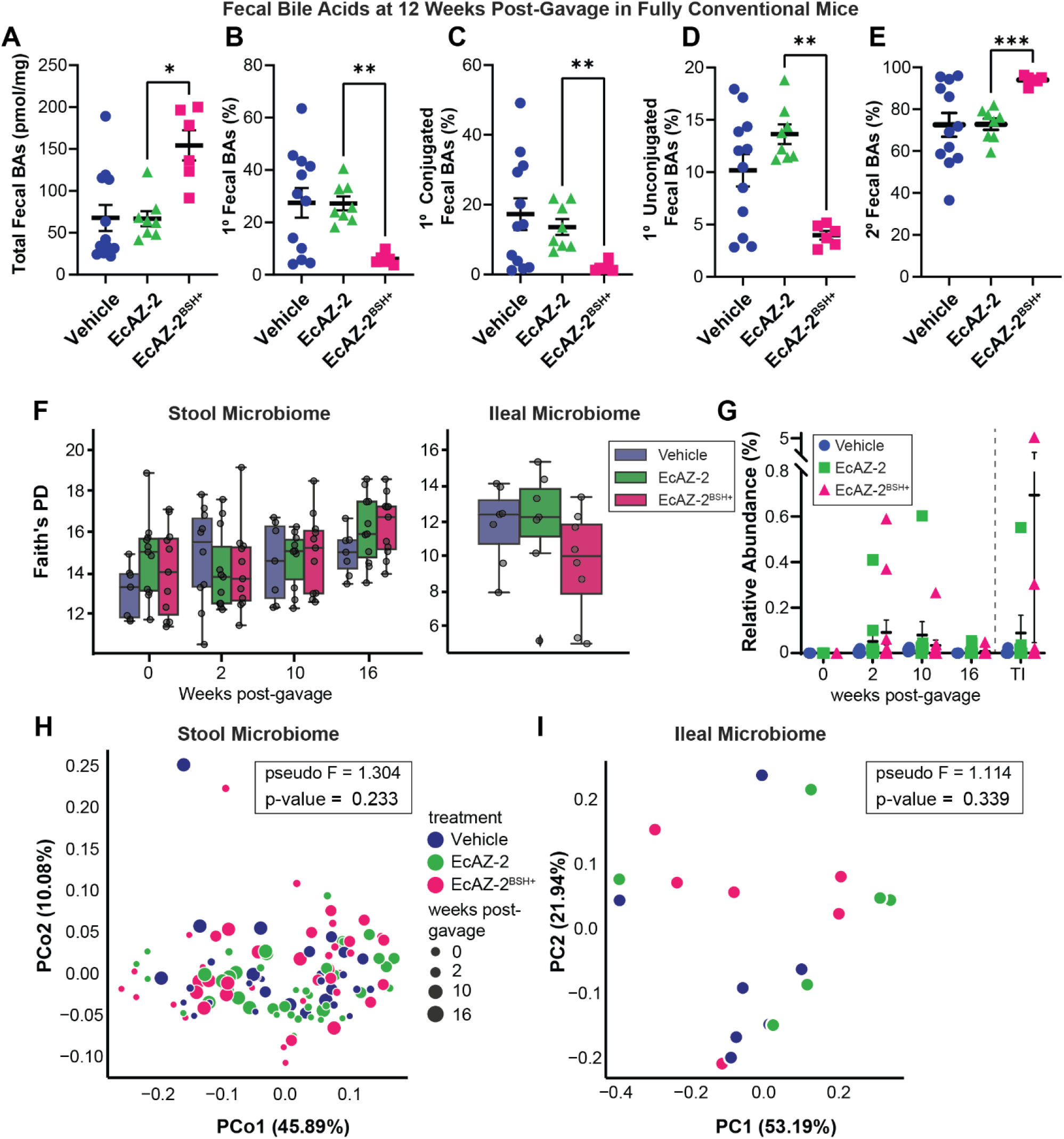
Native *E. coli* can be used to change luminal metabolome without measurable effects in the microbiome. (**A**) Total fecal bile acids, (**B**) Primary bile acids, (**C**) Primary conjugated bile acids, (**D**) Primary unconjugated bile acids, (**E**) And secondary bile acids in fecal samples collected from mice 12 weeks after a single gavage with vehicle, EcAZ-2, and EcAZ-2^BSH+^. Significant differences were determined by Kruskal-Wallis test with post-hoc Dunn’s multiple comparison test comparing EcAZ-2 and EcAZ-2^BSH+^. (**F**) Faith’s phylogenetic distance (a measure of α-diversity) from 16S performed on stool samples collected pre-treatment and at 2-, 10-, and 16-weeks post-gavage (left) and from the terminal ileum samples at the time of euthanasia (right). (**G**) Relative abundance of *E. coli* as detected by 16S performed on stool samples collected pre-treatment and at 2-, 10-, and 16-weeks post-gavage and from the terminal ileum samples at the time of euthanasia. (**H**) Principal coordinate analysis of weighted UniFrac β-diversity of the fecal samples collected pre-treatment and at 2-, 10-, and 16-weeks post-gavage. (**I**) Principal coordinate analysis of weighted UniFrac β-diversity of the terminal ileum microbiome at the time of euthanasia.

We expected that changes in intraluminal bile acids would lead to global changes in gut microbiome composition. However, microbiome analysis showed that EcAZ-2 and EcAZ-2^BSH+^ had no detectable differences in any microbiome measures. Compared to mice treated with vehicle, we found no significant differences in Faith’s phylogenetic diversity in the fecal microbiome at two-, ten-, and sixteen-weeks post-gavage or in the ileum after euthanasia (**Fig. 3F**). Other measures of α-diversity such as richness (**Fig. S3E**) and Shannon index (**Fig. S3F**) found a transient difference at week two between mice in the three conditions both in the fecal or ileal microbiomes that disappeared by the later time points.

There were no *E. coli* detectable by ASV matching the 16S sequence of *E. coli* prior to gavage. Although *E. coli* could be identified in some of the samples from mice treated with EcAZ-2 or EcAZ-2^BSH+^, most had levels of *E. coli* in stool and were statistically indistinguishable as a group from the vehicle treated controls (**Fig. 3G**). Overall, based on 16S sequencing, *E. coli* comprised <0.1% of the fecal microbiome. Principal coordinate analysis of the UniFrac weighted β-diversity performed at two-, ten-, and sixteen-weeks could not distinguish the three treatment groups (**Fig. 3H**; pseudo-F = 1.304, p = 0.233). The lack of a differences is reflected in the ileum as well (**Fig. 3I**; pseudo-F = 1.114, p = 0.339) (**Fig. 3A, 3D**). This demonstrates that, even at low abundances, the native *E. coli* can be used to introduce a function of interest in CR hosts, and they do so without significantly altering the microbial community already established in the host.

### Engineered native E. coli affect host physiology

We next investigated whether transgene delivery and expression by the native *E. coli* chassis can affect host physiology by first determining the impact of colonization with EcAZ-2^BSH+^ on circulating bile acids. Mice colonized with EcAZ-2^BSH+^ trended towards reduced total serum bile acids, increased primary bile acids, and decreased secondary bile acids (**Fig 4A, S4A**). There were larger changes in specific circulating bile acids, including known agonists and antagonists of FXR, such as increased TCA and TbMCA, respectively, compared to their deconjugated counterparts (**Fig. 4B, 4C, S4B**). We measured the impact of these differences in circulating bile acids on hepatic gene expression, focusing on genes important for bile acid metabolism. We found increased expression of *Fxr, Shp*, and *Cyp27a1* (**Fig. 4D**) consistent with increased bile acid biosynthesis through the alternative pathway, perhaps to alleviate the higher excretion of bile acids in the stool of mice colonized with EcAZ-2^BSH+^. This demonstrates that addition of a single gene into the microbiome via engineered native bacteria can significantly alter host physiology.

**Figure 4.**
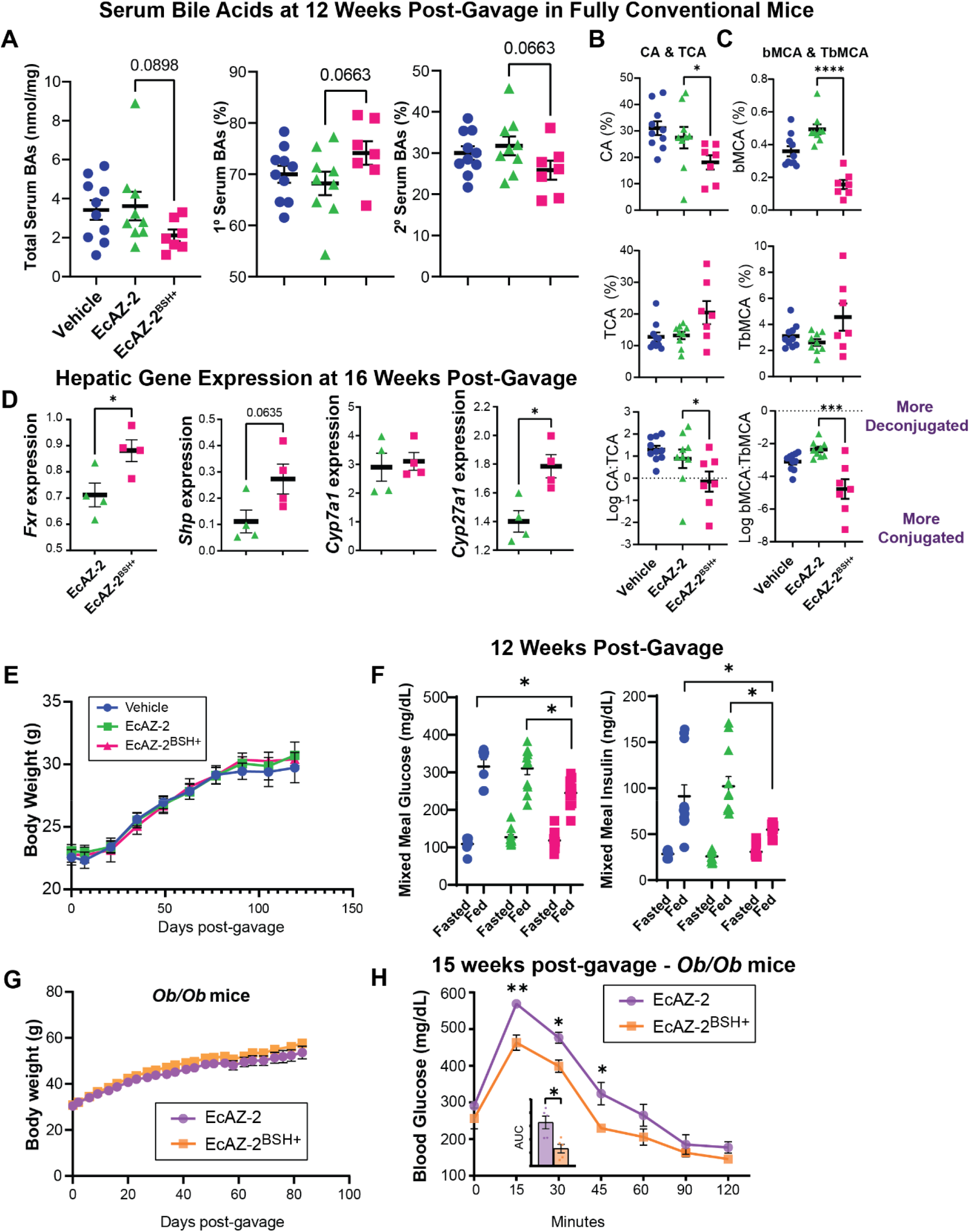
Native *E. coli* can be used to physiologically change the host and treat pathophysiological conditions. (**A**) Total, primary, and secondary serum bile acids from mice treated with a single gavage of vehicle, EcAZ-2 or EcAZ-2^BSH+^ 12 weeks prior. Significant differences were determined by Kruskal-Wallis test with post-hoc Dunn’s multiple comparison test comparing EcAZ-2 and EcAZ-2^BSH+^. (**B**) Serum levels of CA (top), TCA (middle) and the log_2_ ratio of CA to TCA in mice treated with vehicle, EcAZ-2 or EcAZ-2^BSH+^. Significant differences determined by Kruskal-Wallis test with post-hoc Dunn’s multiple comparison test comparing EcAZ-2 and EcAZ-2^BSH+^. (**C**) Serum levels of bMCA (top), TbMCA (middle) and the log_2_ ratio of bMCA to TbMCA in mice treated with vehicle, EcAZ-2 or EcAZ-2^BSH+^. Significant differences determined by Kruskal-Wallis test with post-hoc analysis comparing EcAZ-2 and EcAZ-2^BSH+^. (**D**) Hepatic gene expression of *Fxr, Shp, Cyp7a1*, and *Cyp27a1* as determined by qRT-PCR. Significant differences were determined by a student’s t-test with normality verified through Q-Q plot. (**E**) Mouse weights of CR-WT mice treated with a single gavage of vehicle, EcAZ-2 and EcAZ-2BSH+. (**F**) Fasting (16 hour) and postprandial (30 min) glucose and insulin levels. Significant differences were determined by Kruskal-Wallis test with post-hoc Dunn’s multiple comparison test comparing all three conditions done separately for fasted and fed measures. (**G**) Mouse weights of *Ob/Ob* mice treated with a single gavage of EcAZ-2 and EcAZ-2^BSH+^. (**H**) Glucose tolerance test in *Ob/Ob* mice treated with a single gavage of EcAZ-2 and EcAZ-2^BSH+^ 15 weeks prior. Significance determined with Mann-Whitney U test corrected for multiple comparisons. Inset shows area under the curve. Significance determined by Mann Whitney U test.

To determine whether the transgene delivery with the native *E. coli* chassis can induce long-lasting change in host physiological phenotype, we metabolically characterized the cohort of mice that received vehicle, EcAZ-2, or EcAZ-2^BSH+^ approximately 12 weeks after gavage. Unlike what has been reported in gnotobiotic and reduced community mice,^25, 27^ increased BSH activity induced with engineered native bacteria did not affect the body weight of the host (**Fig. 4E**). The addition of EcAZ-2 or EcAZ-2^BSH+^ did not affect food intake (**Fig. S4C**). Twelve weeks after a single gavage with engineered native bacteria, CR mice on a normal chow diet colonized with EcAZ-2^BSH+^ had reduced blood glucose and insulin levels compared to EcAZ-2 after a mixed meal challenge (**Fig. 4F**). We repeated the mixed meal challenge in a separate experiment using female mice with a shorter duration of colonization with engineered native bacteria (**Fig. S4D**). As with the male mice, the female mice treated with EcAZ-2^BSH+^ had lower postprandial insulin levels **(Fig. S4E)**.

Decreased postprandial insulin induced by EcAZ-2^BSH+^ suggests improved insulin sensitivity. To determine whether transgene delivery with a native *E. coli* chassis could be used to ameliorate disease, we investigated the effect of EcAZ-2^BSH+^ in a genetic model of type 2 diabetes, the *Ob/Ob* mice. EcAZ-2 and EcAZ-2^BSH+^ stably colonized CR *Ob/Ob* mice (**Fig. S4F**). Similar to what we had observed in CR, WT mice, EcAZ-2^BSH+^ did not induce a change in body weight (**Fig. 4G**). Despite identical body weights, *Ob/Ob* mice colonized with EcAZ-2^BSH+^ had significant improvement in insulin sensitivity compared to the control cohort (**Fig. 4H**), after receiving a single treatment of engineered native bacteria more than 3 months earlier. Thus, these experiments show that transgene delivery using native *E. coli* affects the serum metabolome, extra-intestinal (e.g., hepatic) gene expression, host physiology, and ameliorates disease months after a single administration.

### Humans have genetically tractable native E. coli

To determine whether the engineered native bacteria strategy can be translated to humans, we isolated *E. coli* strains from biopsies obtained from multiple regions of the gastrointestinal tract from volunteers undergoing routine outpatient endoscopy. Nine isolated strains were assessed for antibiotic sensitivity, tractability via transduction, transformation with assorted sizes of plasmids, and conjugation (**Table 1**). All but one isolate were sensitive to commonly used laboratory antibiotics including kanamycin, chloramphenicol, and carbenicillin. All isolates evaluated were resistant to transduction via P1*vir* when lysates were generated from *E. coli* MG1655 or *E. coli* Nissle 1917. Each strain was transformable with small plasmids (roughly 7 kb), but not larger plasmids (>30 kb). Conjugation using a modified F-plasmid proved to be a universally successful method for manipulation. To encourage plasmid retention, conjugation machinery was removed post-transfer. Thus, the first steps of the engineered native bacteria approach, identification/isolation of genetically tractable bacteria and modification to express a gene of interest, can be translated to humans. Further studies are necessary to determine whether the autologous transfer of engineered native bacteria can lead to long-lasting colonization of a human host and treatment of long-standing chronic conditions and genetic diseases.

**Table 1:**
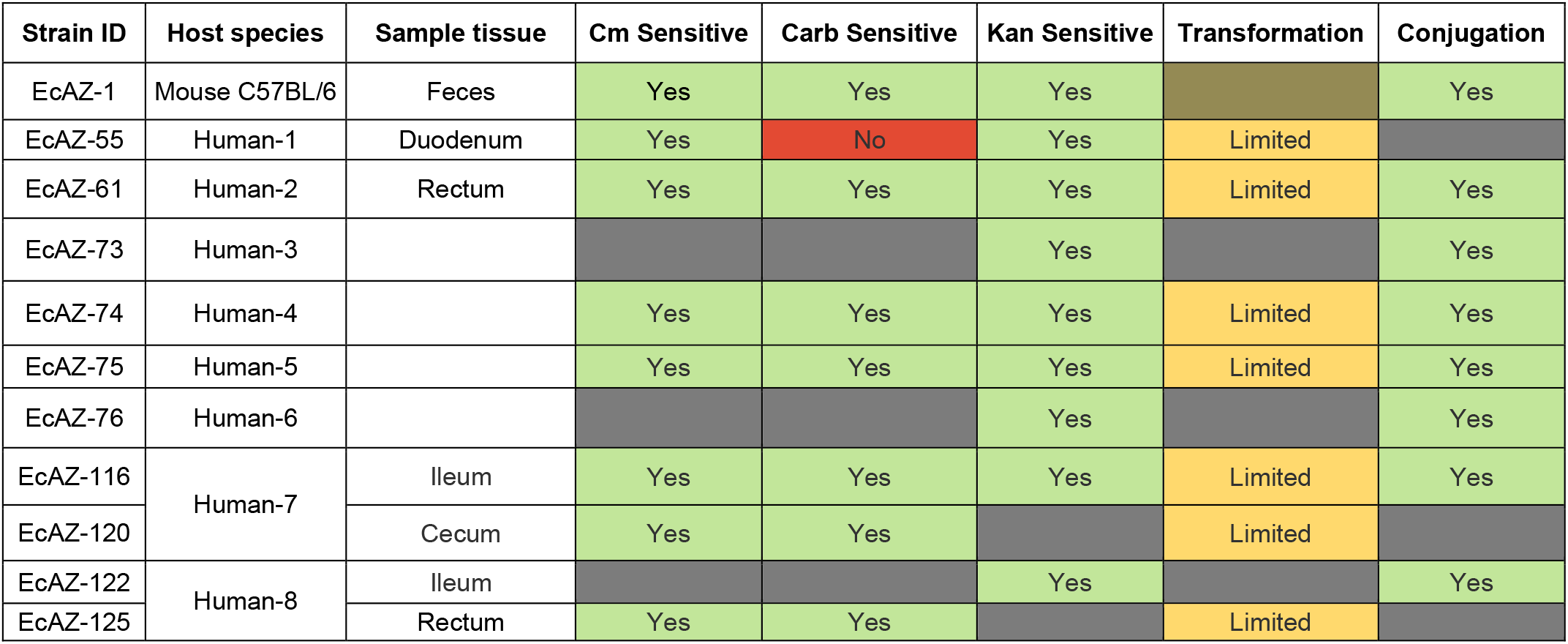
Human derived *E. coli* isolated from intestinal biopsies are genetically tractable.

## DISCUSSION

Most therapies that target microbiome composition do not have a detectable impact on the gut microbiome and are not robust to the interpersonal diversity and plasticity of the microbiome in human hosts.^15, 30^ In addition, many different configurations of the gut microbiota lead to the same functional result,^31, 32^ suggesting that microbial function may be difficult to manipulate with therapies that target the composition of the microbiome alone. To develop a better mechanistic understanding of the microbe-host relationship and more effective microbiome-mediated therapies, approaches based on functional modulation of the gut microbiome are necessary. However, these approaches have been difficult to develop, and none have demonstrated long-term engraftment with a change in physiology and/or improvement of pathologic phenotype.

To address the inability of engineered bacteria to colonize the CR host, only two strategies have shown success:^14, 15, 33-35^ creating exclusive metabolic niches for an engineered probiotic strain which can then outcompete resident microorganisms,^36^ and engineering non-colonizing bacteria to distribute mobile genetic elements (e.g., plasmids) to native bacteria *in vivo*.^37^ However, neither approach has been demonstrated to alter luminal or serum metabolome, affect host physiology, or demonstratively ameliorate disease as shown here.

A promising approach has been to find genes that improve engraftment in commonly used chassis, such as *E. coli* Nissle 1917.^38^ More commonly, investigators have been developing tools to manipulate families of bacteria that are found in large numbers in the gut microbiome to determine whether they can outcompete or coexist with native strains and engraft in the intestinal lumen.^39-42^ These bacteria and other engineered probiotics have been exclusively assessed under reduced community (as opposed to CR) conditions (i.e., gnotobiotic mice, antibiotic-treated mice). These efforts have advanced our understanding of bacterial engraftment, biogeography of microorganisms and luminal micro-niches, and the biological requirements for bacterial colonization and function delivery. However, they also indicate that the amount of bacterial engineering and experimentation necessary to develop chassis that can colonize the gut is daunting and out of the reach for resource-limited labs without dedicated synthetic biologists. Moreover, they are problematic for studying steady state mechanisms in the gut, unlikely to work in animal models where diet and host genetics are manipulated, and limit feasibility of these techniques for adoption and interventions in humans.

Here we demonstrate a simple technique that advances our ability to use engineered bacteria to effectively change physiology in CR-WT hosts, using tools and approaches that can be adopted by nearly any lab. By combining existing techniques to native strains, we demonstrate that undomesticated *E. coli* derived from the host can be used as chassis to knock in specific functions into the gut microbiome, thus making mechanistic studies of the gut microbiome attainable to most labs without advanced synthetic biology experience. *E. coli* are easily culturable, common in the human gut,^43-47^ and tools to engineer them have existed for decades. Although *E. coli* are common in the gut lumen, many have assumed that they are not good colonizers due to so much disappointment with commensal lab strains, such as *E. coli* Nissle. However, we show that by using native *E. coli* as our chassis, the resultant engineered native bacteria can (a) stably colonize CR-WT hosts for months, possibly in perpetuity after a single gavage, without the need for pretreatment for microbiome depletion, (b) engraft the proximal gut in addition to the colon, (c) demonstrate natural biocontainment between co-housed mice, (d) alter specific luminal metabolites as intended, that lead to (e) a change in serum metabolites, which then (f) change host metabolic homeostasis and core physiological processes, and, thus, (g) prevent dysmetabolic phenotypes. Moreover, our preliminary data show that the initial steps of this process can be translated to humans, where we have found readily tractable, native *E. coli* which could be transformed to express a gene. Importantly, we show that identical cohorts who only differ in a single gene in a single bacterial strain can have a dramatically different metabolic phenotype. Although this has been demonstrated with pathogenicity islands (e.g., Shiga toxin), we now demonstrate that a beneficial bacterial gene can also have systemic effects.

Earlier studies of an *E. coli* strain isolated from mouse fecal samples from a different facility (i.e., NGF-1)^48^ likewise demonstrated long-term colonization.^21^ However, it was not clear whether undomesticated *E. coli* strains could be used to study specific bacterial functions, affect the luminal environment, and shape host physiology. Given that *E. coli* abundance is < 0.1% of the microbiome, others have focused on chassis that belong to more highly abundant strains. Interestingly, our study shows that colonization of the host luminal environment with our engineered *E. coli* did not affect the overall composition of the microbiome, nor the relative abundance of *E. coli* as detected by 16S analysis. This suggests several possibilities including that a) *E. coli* potentially occupies a particular niche that enables close interaction with the host without too much change in the overall composition of the microbiome, and b) *E. coli* is replacing or co-existing with its clonal relatives. Large microbiome resource studies, such as the Human Microbiome Project and American Gut Project,^49, 50^ show limited detection of *E. coli* by 16S sequencing from fecal samples, whereas infants have higher levels,^51^ suggesting that *E. coli* abundance may be affected by niche availability. Moreover, low abundance individual microbes can have large ecosystem effects because of the niche they occupy and functions they perform;^52^ thus, a microbe need not be abundant to have a big impact. Overall, our ability to find genetically tractable *E. coli* in humans shows that this approach can be translatable in a vast majority of patients.

Our results clearly demonstrate that native *E. coli* are particularly adept at affecting host physiology and, considering the engineering toolbox that has been developed to genetically manipulate *E. coli*, make an effective chassis. In addition, having chassis that are unable to engraft has been proposed as a mechanism for biocontainment since it would improve the safety profile by removing concern for LBT proliferation, transmission to healthy cohabitants, and therapy delivery beyond the end of treatment.^12, 14^ However, given the inability of current developed methods to affect change in CR hosts, including humans, the development of techniques to make transgene expression tunable,^41, 42^ and our results showing biocontainment with cohoused mice, we suggest that transgene delivery with engineered native chassis will be a far more effective approach.

The technical capability to change microbiome function in CR mice is critical to understanding the role of the gut microbiome in various physiological and pathophysiological processes. Many animal models of disease cannot be recapitulated in reduced microbiome community conditions. For example, baseline metabolic activity in microbiome-depleted (e.g., gnotobiotic, antibiotic treated) mice are vastly different from conventionally-raised mice.^53, 54^ This could be why the effects of BSH on host metabolic physiology have been difficult to understand, since prior studies using different reduced community models have led to contradictory results.^25, 27^ Because deconjugated bile acids do not have dedicated transporters and require reabsorption through passive diffusion, increased luminal BSH activity should lead to an increase in fecal excretion of bile acids, though this has not been reported in microbiome depleted models.^25^ Moreover, they ignore the downstream compositional, metabolomic, and proteomic effects that can lead to changes to the luminal community that could amplify or dampen the effects of the knocked-in function. For example, though increased BSH activity is hypothesized to lead to an increase in primary unconjugated bile acids, we observed the opposite. Instead, because deconjugated bile acids are substrates that some bacteria can convert to secondary bile acids, we observed an increase in secondary bile acids. This observation is apparent in CR mice but not in mono-colonized or reduced-community mice. Although our present work focuses on providing a proof-of-concept study, the BSH results show something different from reduced-community studies - a decrease in serum insulin in the setting of unchanged body weight - which profoundly changes our understanding of how microbiome bile acid biotransformations affect host metabolism.^55^ Because human metagenomic studies have identified BSH as protective against diabetes,^56, 57^ additional studies in the CR hosts are necessary to better understand the effect of bile acids on host metabolism and other physiological processes.

The strategy of using native bacteria as a chassis to introduce transgenes of interest can be used for different organ systems (e.g., skin, lungs, vagina), expanding our ability to perform mechanistic studies and providing a strategy for the creation of therapeutic engineered native bacteria in any microbiome mediated/modulated disease. Moreover, personalized engineered native bacteria will help overcome the barriers that have limited the use of engineered bacteria in humans, including interpersonal differences and micro-niche changes induced by disease. Finally, life-long compliance with chronic therapies is a substantial burden in patients with a number of chronic diseases, which contributes to their poor morbidity and mortality, and to rising healthcare costs. Some engineered probiotics in development for disease intervention require multiple doses (sometimes up to three times daily), which may add significant complications for feasibility, cost, and adherence. If the engineered native bacteria strategy is successfully developed and translated to humans, it has the potential to introduce novel, curative biotherapeutics that improve treatment of chronic diseases without relying on patient compliance.

## ACKNOWLEDGEMENTS

We would like to express our appreciation to Dr. Kim Barrett and Dr. David Brenner, who provided resources to initiate this project during its conceptualization. SDB was supported by F32 DK113721. ARS is supported by the Glenn Foundation for Medical Research Postdoctoral Fellowships in Aging Research. IM was supported by a UCSD Eureka Foundation Grant. NS is a Biolegend Fellow. EM is supported by F31 HD106762. ACDM is supported by R01 HL148801-02S1. AZ is supported by AFAR Research Grant for Junior Faculty, American Heart Association Beginning Grant-in-Aid (16BGIA27760160), Kavli Institute for Brain and Mind at UC San Diego, Jon I. Isenberg Endowed Fellowship, AASLD Liver Scholar Award, AGA Microbiome Junior Investigator Award, and NIH K08 DK102902, R03 DK114536, R21 MH117780, R01 HL148801, R01 EB030134, and U01 CA265719. All authors receive institutional support from NIH P30 DK120515, P30 DK063491, P30 CA014195, P50 AA011999, and UL1 TR001442. The funders had no role in study design, data collection and interpretation, or the decision to submit the work for publication.

## CONTRIBUTIONS

Conceptualization: SDB, BJR, AZ; Methodology, Investigation, and Validation: SDB, BJR, ARS, IM, AL, NS, EM, YM, AFMP, AZ; Software: ACDM, RAR; Formal Analysis: BJR, ACDM, RAR, AZ; Resources: SBH, LE, AS, RK, AZ; Data Curation: RAR; Writing – Original Draft Preparation: BJR, AZ; Writing – Reviewing and Editing: BJR, SDB, RAR, SBH, LE, JH, AS, RK, AZ; Visualization: BJR, AZ; Supervision: AZ; Project administration: AZ; Funding Acquisition: AZ.

## DATA AND MATERIAL AVAILABILITY

EcAZ-1, EcAZ-2, EcAZ-2^BSH+^ and EcAZ-2^IL10+^ will be made available subject to a materials transfer agreement with the University of California, San Diego. Any and all data and code relating to the paper will be deposited in a public database with a release date set to the date of publication.

## MATERIALS AND METHODS

### Bacterial Isolation

For murine derived strains, stool was collected from a male C57BL/6 male mouse acquired from Jackson Laboratory (Bar Harbor, ME). Stool was suspended in sterile DI water and homogenized for 2 min. at 3500 rpm in Mini-Beadbeater-24 (Biospec, Bartlesville, OK). Homogenized stool was plated on MacConkey agar containing lactose. Potential *E. coli* colonies were selected based on colony shape and ability to ferment lactose. They were cultured for isolation twice. 16S rRNA genes were amplified using the primers described below (Illumina Part #15044223 Rev. B) and sequenced using Sanger sequencing (Eton Biosciences, La Jolla, CA) to confirm identity.

For human-derived strains, volunteers undergoing outpatient endoscopy were selected in accordance with VA San Diego IRB #H130266. Biopsies were collected from various regions of the GI tract adjacent to sites that were needed for clinical indications (e.g., polyps) and immediately transferred to sterile PBS. Tissue samples were homogenized, and bacterial growth, isolation, and identification was performed as described above. 16S rRNA gene amplicon PCR forward primer =

5′-TCGTCGGCAGCGTCAGATGTGTATAAGAGACAGCCTACGGGNGGCWGCAG

16S amplicon PCR reverse primer =

5′-GTCTCGTGGGCTCGGAGATGTGTATAAGAGACAGGACTACHVGGGTATCTAATCC

### Strain Development

The index native murine *E. coli* used for transformation was labeled as EcAZ-1. For EcAZ-2, ECAZ-1 was exposed to P1*vir* lysate trained on MG1655 with kanamycin resistance (*aph*) and fluorescence (*GFP-mut2*). Successful transductants were selected for using kanamycin resistance (*aph*) and fluorescence (EcAZ-2). EcAZ-1 was nearly 100-fold more resistant to phage transduction than *E. coli* Nissle. For the construction of EcAZ-2 containing bile salt hydrolase (BSH; EcAZ-2^BSH+^), EcAZ-1 was exposed to P1*vir* lysate trained on MG1655 containing chloramphenicol resistance (*cat*) and *bsh*. Successful transductants were selected for functionality of BSH and chloramphenicol resistance. Verification of BSH functionality was verified by plating on lactose-containing LB plates supplemented with TDCA. Precipitation of DCA indicated functioning BSH. The successful transductants were then exposed to lysate with kanamycin resistance (*aph*) and fluorescence (*GFP-mut2*). Successful transductants were selected for gain of kanamycin resistance. The bsh gene used for this study naturally occurs in *Lactobacillus salivarius* JCM1046 and is effective at altering bile acids in germ free and antibiotic-treated, conventionalized mice when expressed by an *E. coli* Nissle chassis.^25^ The first insertion event (for the cassette containing *aph* and *gfp*) occurred between chromosomal bases 3,267,361-3,274,418, which covers a Ribosomal RNA operon, *murI* (glutamate racemase) and *btuB* (vitamin B12 / E colicin / bacteriophage BF23 outer membrane porin BtuB). The second insertion event (for the cassette containing *cat* and *bsh*) occurred between chromosomal bases 1,886,038-1,887,917, which covers *ydcM* (putative transposase), *ybhB* (putative kinase inhibitor), and *ybhC* (outer membrane lipoprotein).

For the construction of EcAZ-2 containing IL-10 (EcAZ-2^IL10+^), EcAZ-1 was exposed to a lysate trained on MG1655 containing chloramphenicol resistance (*cat*) and human *Il10*. The successful transductants were selected for via IL-10 gene detection by PCR and chloramphenicol resistance. Insertion occurred in the *yfgG* (metal ion stress response protein) gene. Verification of IL-10 expression was performed by growing an overnight culture of isolates in Super Optimal Broth (SOB), 12.5 ug/mL of Kanamycin and 10 ug/mL of chloramphenicol. The cultures were centrifuged and resuspended in Assay diluent (10% FBS in 1x PBS). The samples underwent four freeze-thaw cycles, and the cell lysates were assessed on a human IL-10 detection ELISA (Catalog 430601, Biolegend).

Human bacterial isolates were assessed for susceptibility to commonly used laboratory antibiotics at standard laboratory concentrations (10 ug/mL for chloramphenicol, 12.5 ug/mL and 25 ug/mL for kanamycin, and 100 ug/mL for carbenicillin). Human strains were assessed for susceptibility to transduction via P1*vir* using lysate trained on MG1655 and lysate trained on *E. coli* Nissle 1917. Transformation was performed via electroporation. Three plasmid sizes were assessed including pTrcHisB, a modified version of pNCM (roughly 70kb), and an additionally modified version of pNCM (roughly 35kb). All plasmid transformations were assessed at 1800 V, 200 ohms, and 25 uF. Conjugation was performed using a 2,6-diaminopimelic acid dependent donor strain containing a modified F-plasmid, pNCM, containing a kanamycin resistance marker and green fluorescent protein with a deletion in *finO* to promote conjugation. Successful conjugates were selected for the ability to grow without DAP, and for a gain of GFP and kanamycin resistance.

### Growth Curves

Overnight cultures of SOB, supplemented with no antibiotic for EcAZ-1, 12.5 ug/mL of kanamycin for EcAZ-2, and 10 ug/mL of chloramphenicol for EcAZ-2^BSH+^ and EcAZ-2^IL10+^, were back diluted 1:100 in the same media described and loaded in triplicate in a 96-well plate. Blanks for each media were also loaded in triplicate. Incubation and measurements were done in an Infinite M Nano+ plate reader (Tecan, Morrisville, NC) at 37°C. OD600 measurements were performed every 10 minutes for 18 hours directly after 5 seconds of shaking. OD was determined by subtracting out the average blank for the media type for each time point.

### Animal Experiments

All animal experiments were conducted in accordance with the guidelines of the IACUC of the University of California, San Diego. C57BL/6 mice (Jackson Laboratories) were housed in a specific pathogen free facility (irradiated chow and autoclaved bedding), or when specified, in a separate section of the vivarium that was a low barrier facility (non-irradiated chow and non-autoclaved bedding; e.g., **Fig. 1A, D**). Mice were housed 4-5 per cage unless otherwise stated. They were fed a normal-chow diet (Diet 7912, Teklad Diets, Madison, WI). After an acclimation period, 12-week-old male mice were pseudorandomized into two groups. This pseudo-randomization was based on the initial weight of the mice; in the end, the mean and standard deviation of the weight of the mice was the same between all groups. Mice were gavaged with 200 µL of either PBS or ∼1×10^10^ cfu/mL of the corresponding bacterial strain. Body weight and food consumption were monitored every 1-2 weeks.

### Colonization Assessment

For stool, we collected 3-5 stool pellets from individual mice and weighed them. 1 mL of sterile deionized water was added to each sample along with one sterile chrome bead (Neta Scientific, Hainesport, NJ). Samples were homogenized for 2 min. at 3500 rpm in Mini-Beadbeater-24 (Biospec, Bartlesville, OK). For control samples, homogenized stool was plated on kanamycin and chloramphenicol to check for bacterial contamination. For EcAZ-2 colonized mice, samples were plated on chloramphenicol to check for unintended bacterial contamination and diluted and plated on kanamycin to calculate CFU/gram of stool. The EcAZ-2^BSH+^ and EcAZ-2^IL10+^ stool was diluted and plated on kanamycin and chloramphenicol.

Collected and tested tissues include duodenum defined as the first 10 cm of small intestine, jejunum defined as the next 10 cm of small intestine, Ileum defined as the next 5 cm of small intestine, cecum, and colon.

For tissues, mice were euthanized, and tissue samples were harvested and immediately placed on ice. Dissection tools were disinfected between animals. Samples were processed and plated as described for stool. For each region of the GI tract, bacterial colonies were evaluated for enzyme functionality by patching from non-selective plates onto lactose and taurodeoxycholic acid (TDCA) containing plates. Up to 25 colonies for each region for each mouse were evaluated. In cases where 25 colonies did not grow, all colonies were evaluated.

For each region of the GI tract, verification of IL-10 expression was performed by growing an overnight culture of isolates in SOB, 12.5 ug/mL of Kanamycin and 10 ug/mL of chloramphenicol. To lyse the bacteria, the cultures were centrifuged, and the resulting pellet was resuspended in Assay diluent (10% FBS in 1x PBS). The samples were then flash frozen and defrosted for 4 rounds to lyse the cells and used on an IL-10 detection ELISA (Catalog 430601, Biolegend).

### Co-Housing Experiment

Three pairs of 6-month-old male C57BL/6 mice were co-housed in a low barrier facility (where food and bedding were not autoclaved). From each cage, one mouse was gavaged with 200 µL of PBS while the other was gavaged with 200 µL ∼1×1010 cfu/mL EcAZ-2. After gavage, mice were single housed for a week. Five days post-gavage, baseline stool was collected from each mouse. The pairs were then reunited with their cage mate. Initial stool collection demonstrated coprophagia but no persistent colonization. After 19 days, the protocol was modified so that mice were separated and individually housed one day prior to stool collection. After stool collection, each mouse was reunited with their cage mates until the next stool collection. Thus, with the new protocol, coprophagic incidents would no longer appear and only incidents of horizontal transfer/colonization would be detectable.

### Gnotobiotic Mice

Germ-free C57BL/6 mice were bred and maintained in sterile flexible film isolators and screened for bacterial, viral, and fungal contamination as described.^58^ Mice were fed autoclaved chow (Envigo 2019S) and transferred into the Sentry SPP cage (Allentown). Mice were inoculated with corresponding bacteria oral gavage. Stool was collected for bile acid analysis 7 days later. Mono-colonization was confirmed upon euthanasia of the gnotobiotic mice.

### Stool and Serum Bile Acids

Bile acids were extracted from samples as described before.^53, 59^ Briefly, stool samples were homogenized and extracted in methanol (10 mg of sample/100 µL) containing heavy internal standards. Serum samples (25 µL) were extracted with 75 µL of methanol containing heavy internal standards. After vortexing for 10 minutes and centrifuging (16,000 x g, 4 C, 10 min), supernatants were transferred to glass vials for injection. Bile acids were analyzed on a Dionex Ultimate 3000 LC system (Thermo) coupled to a TSQ Quantiva mass spectrometer (Thermo) fitted with a Kinetex C18 reversed phase column (2.6 µm, 150 × 2.1 mm i.d., Phenomenex). The following LC solvents were used: solution A, 0.1 % formic acid and 20 mM ammonium acetate in water, solution B, acetonitrile/methanol (3/1, v/v) containing 0.1 % formic acid and 20 mM ammonium acetate. The following reversed phase gradient was utilized: at a flow rate of 0.2 mL/min with a gradient consisting of 25-29 % B in 1 min, 29-33 % B in 14 min, 33-70 % B in 15 min, up to 100 % B in 1 min, 100 % B for 9 min and re-equilibrated to 25 % B for 10 min, for a total run time of 50 min. The injection volume for all samples was 10 µL, the column oven temperature was set to 50°C and the autosampler kept at 4°C. MS analyses were performed using electrospray ionization in positive and negative ion modes, with spray voltages of 3.5 and - 3 kV, respectively, ion transfer tube temperature of 325°C, and vaporizer temperature of 275°C. Multiple reaction monitoring (MRM) was performed by using mass transitions between specific parent ions into corresponding fragment ions for each analyte. The transitions and retention time for each analyte and internal standards are shown in **Table S1**. Results were quantified using isotopically labeled internal standards. Data were averaged across samples.

The bile acids measured using this technique include CA, TCA, bMCA, TbMCA, α- muricholic acid (aMCA), tauro-α-muricholic acid (TaMCA), chenodeoxycholic acid (CDCA), taurochenodeoxycholic acid (TCDCA), DCA, TDCA, hyocholic acid (HCA), taurohyocholic acid (THCA), ω-muricholic acid (oMCA), tauro-ω-muricholic acid (ToMCA), ursodeoxycholic acid (UDCA) and lithocholic acid (LCA). Glycine conjugated bile acids were not detected. Of note, no UDCA was detected in the stool. Total bile acid measurements were the sum of all bile acids measured using mass spectrometry. Primary bile acids included CA, TCA, aMCA, TaMCA, bMCA, TbMCA, CDCA, and TCDCA. Secondary bile acids included DCA, TDCA, oMCA, ToMCA, UDCA, and LCA.

### 16S rRNA Microbiome Data Processing and Analysis

16S rRNA gene amplicon sequences were processed using Qiime2 (version 2020.6) using default parameters.^60^ ASVs were generated by the Deblur method, trimmed to 120bp.^61^ Taxonomy was assigned via the sklearn plugin^62, 63^ against the Silva v123 model,^64^ and ASVs were filtered to remove Eukaryotic, Mitochondrial, Plastid, and unassigned taxa. The phylogenetic diversity metrics and PCoA were then calculated on samples rarefied to 4000 reads.

### qPCR from Liver and Terminal Ileum

The livers and terminal ileum from control and EcAZ-2^BSH+^ mice were flash frozen in liquid nitrogen. The frozen tissues were homogenized, and RNA was extracted using TriZol reagent according to manufacturer’s instructions. 1ug of RNA was reverse transcribed using quanta biosciences qScript cDNA supermix (Cat# 95048-025). The samples were diluted 1:50 and qRT-PCR was run using FastStart Universal SYBR Green Master (Rox) (Cat# 4913850001) on an Applied Biosystems QuantStudio 5. The data were analyzed, and gene expression was expressed relative to mrpl46.

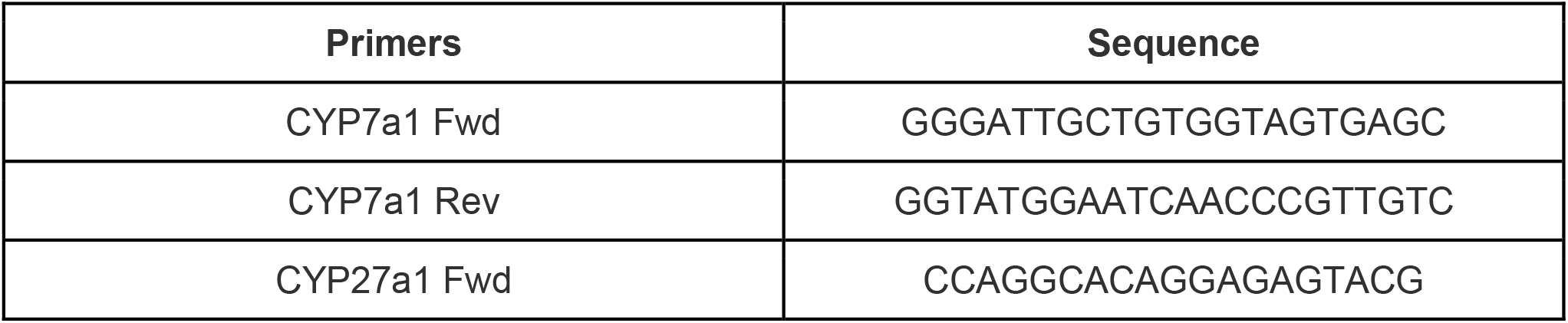
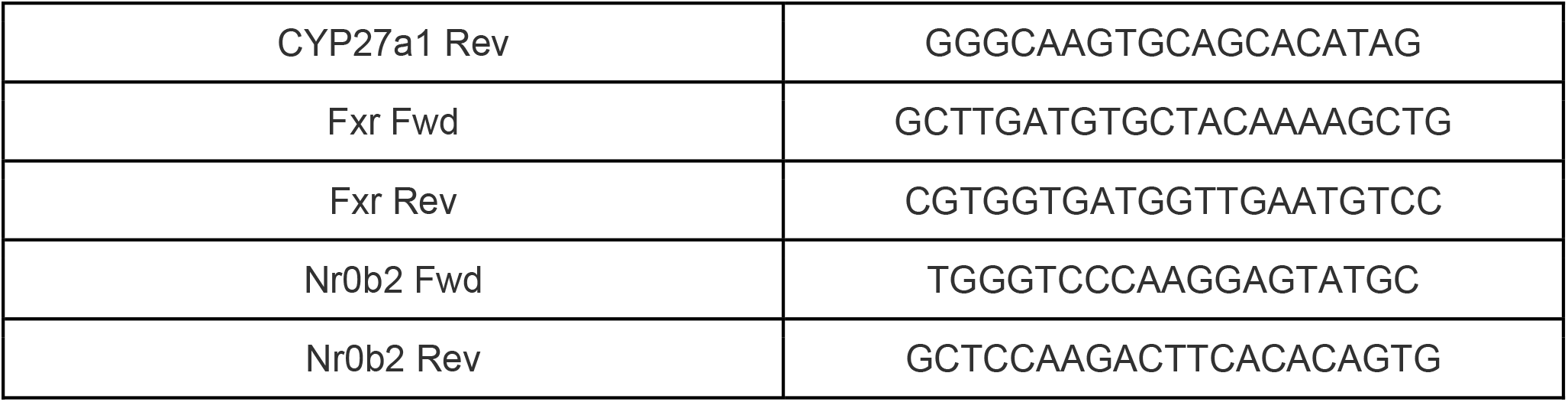

### Blood Collection and Serum Biomarker Measurements

Blood was collected from fasted (16 h) or re-fed animals (30 min). Re-feeding was performed by oral gavage of Ensure (Abbott, Columbus, Ohio). In male mice, serum insulin was performed using the Bio-Plex Pro Mouse Diabetes 8-Plex Assay (Bio-Rad, Hercules, CA) as per the manufacturer’s protocol.

### Glucose Tolerance

Glucose tolerance assays were performed on fasted mice (16 h, Pur-O-Cel bedding) by monitoring glucose levels after a glucose bolus (10 uL of 20% glucose/kg of body weight (BW)) via oral gavage. Glucose concentration was measured from a tail snip using a OneTouch Ultra glucometer. Tail tips were anesthetized with a topical anesthetic (Actavis, Parsippany-Troy Hills, NJ) prior to snip.

### Insulin Measurement

10-week-old female mice were fasted (5 h, Pur-O-Cel bedding) and then bled from the tail vein to collect fasted plasma. Two days later, mice were again fasted (14-16 h, Pur-O-Cel bedding) and then orally gavaged with 0.5g glucose/kg body weight of Ensure (Abbott, Columbus, Ohio) to measure insulin response to a mixed meal. 30 minutes after the gavage, blood was collected through a submandibular bleed. Serum insulin from fasted and fed blood was measured using a mouse ELISA (Crystal Chem Ultra-Sensitive Mouse ELISA Kit).

### Statistics

All comparisons of EcAZ-2 to EcAZ-2^BSH+^ were calculated using Mann-Whitney U test or Kruskal-Wallis test with post-hoc Dunn’s multiple comparison test as specified except for **Fig. S3A-C** and **Fig. 4D**, which used a student’s t-test due to low sample size. When student’s t-test was used, normality of data was confirmed with a Q-Q plot.

