## Supplemental Figures for "Intestinal Transgene Delivery with Native *E. coli* Chassis Allows Persistent Physiological Changes"

********SUPPLEMENTARY MATERIAL********

**<u>Content:</u>**

Figure S1-4

#### Supplemental Figure 1

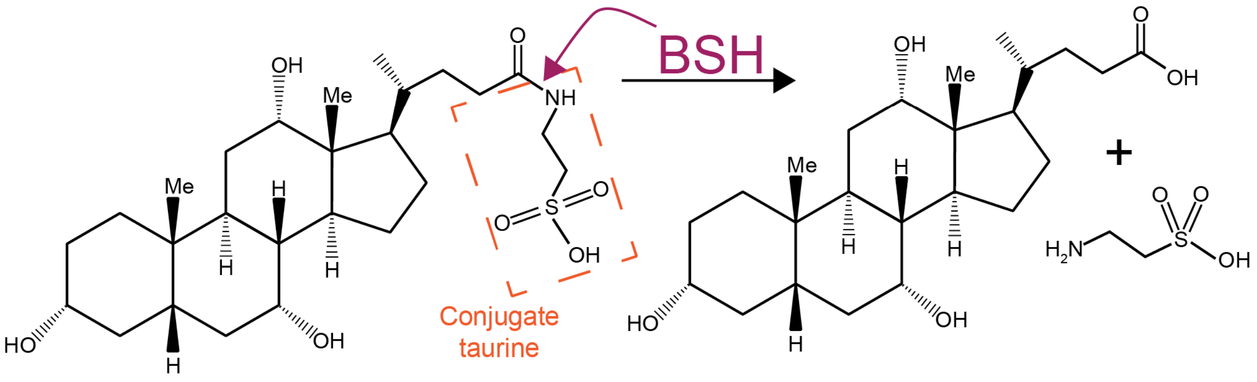


**Supplemental Figure 1 -** Bile salt hydrolase (BSH) is a prokaryotic gene that deconjugates bile acids**.** Through deconjugation, BSH makes bile acids less polar (and thus unable to be transported by apical sodium-bile acid transporters). The deconjugated bile acids can serve as metabolic substrates for a number of bacteria that then convert them to secondary bile acids.

### Supplemental Figure 2

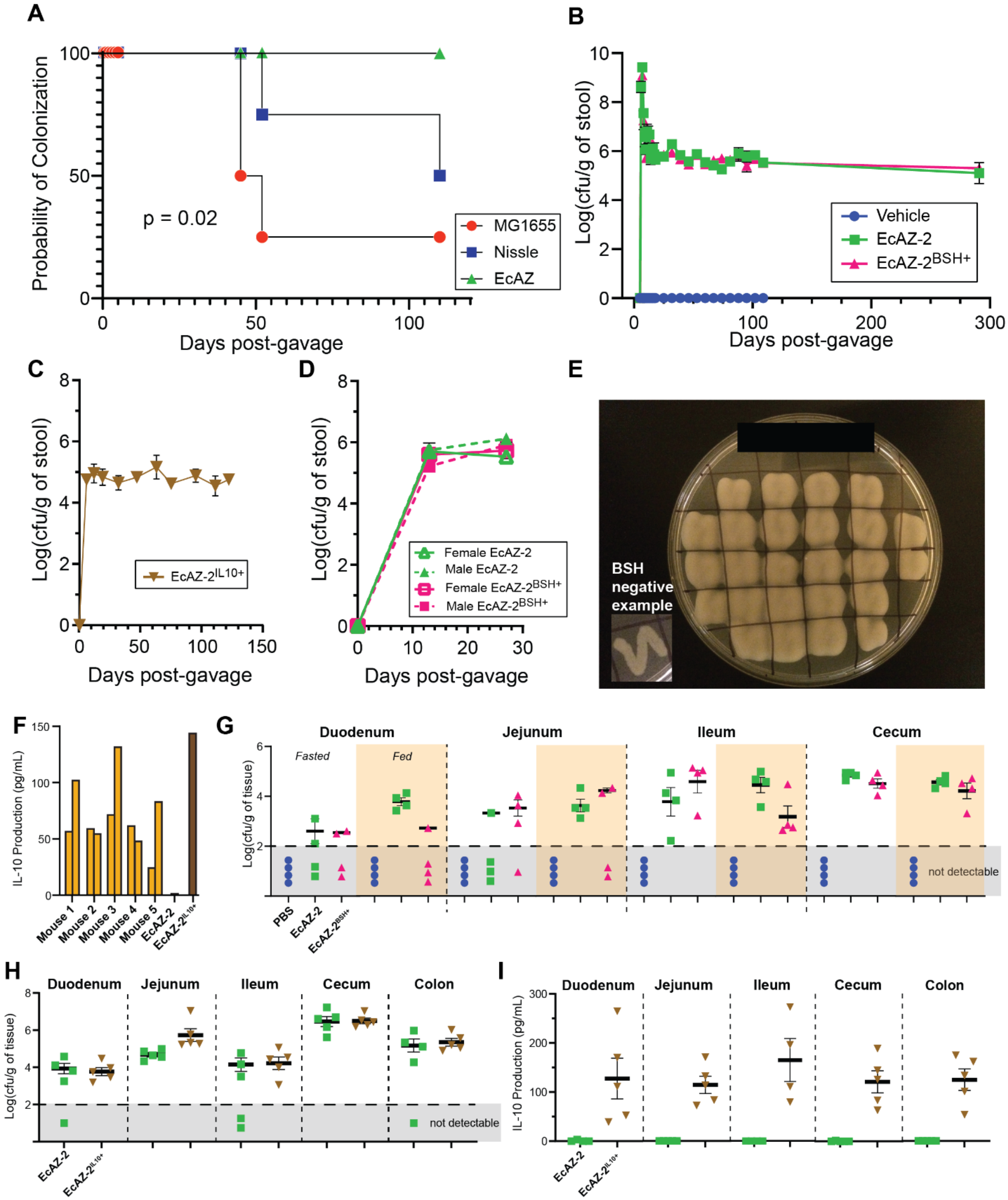


**Supplementary Figure 2**

**(A)** Percentage of mice colonized with their respective bacteria in a non-sterile, low barrier mouse facility (4 mice/condition).

**(B)** Colonization of EcAZ-2 and EcAZ-$2^{BSH+}$ in CR-WT mice housed in a specific-pathogen free facility (4-12 mice/condition).

**(C)** Colonization of EcAZ-$2^{IL10+}$ in CR-WT mice housed in a specific-pathogen free facility (5 mice).

**(D)** Colonization of EcAZ-2 and EcAZ-$2^{BSH+}$ in CR-WT male and female mice housed in a specific-pathogen free facility (4 mice/condition).

**(E)** Example of post-euthanasia testing of colonies from ileum of EcAZ-$2^{BSH+}$ mouse plated on LB containing lactose and TDCA. These strains were isolated 112 days post-gavage from mice who received a single gavage of EcAZ-$2^{BSH+}$. Precipitate around EcAZ-$2^{BSH+}$ is the deconjugated form of TDCA (DCA) and indicates BSH functionality.

**(F)** Two isolates were taken from EcAZ-$2^{IL10+}$ colonies found in the feces of mice depicted in (**C**) from the 89 days post-gavage time point. IL-10 in the cell lysate of these isolates are depicted here. EcAZ-2 is the negative control, whereas EcAZ-$2^{IL10+}$ depicts IL-10 measured from a strain that was never gavaged into a mouse (positive control).

**(G)** Colonization of EcAZ-2 and EcAZ-$2^{BSH+}$ in the gastrointestinal tract of CR-WT mice housed in a specific-pathogen free facility. Measurements were made from mice euthanized after a 16-hr fast, or 1 hour after refeeding. (4 mice/condition).

**(H)** Colonization of EcAZ-2 and EcAZ-$2^{IL10+}$ in the gastrointestinal tract of CR-WT mice housed in a specific-pathogen free facility 79 days after gavage (same mice depicted in **Fig. 2E**; 5/condition).

**(I)** IL-10 levels in the cell lysates of bacteria isolates from mice described in (**H**).

All error bars indicate standard error of the mean. The marker covers some error bars in panel **B**, **C**, and **D**.

### Supplemental Figure 3

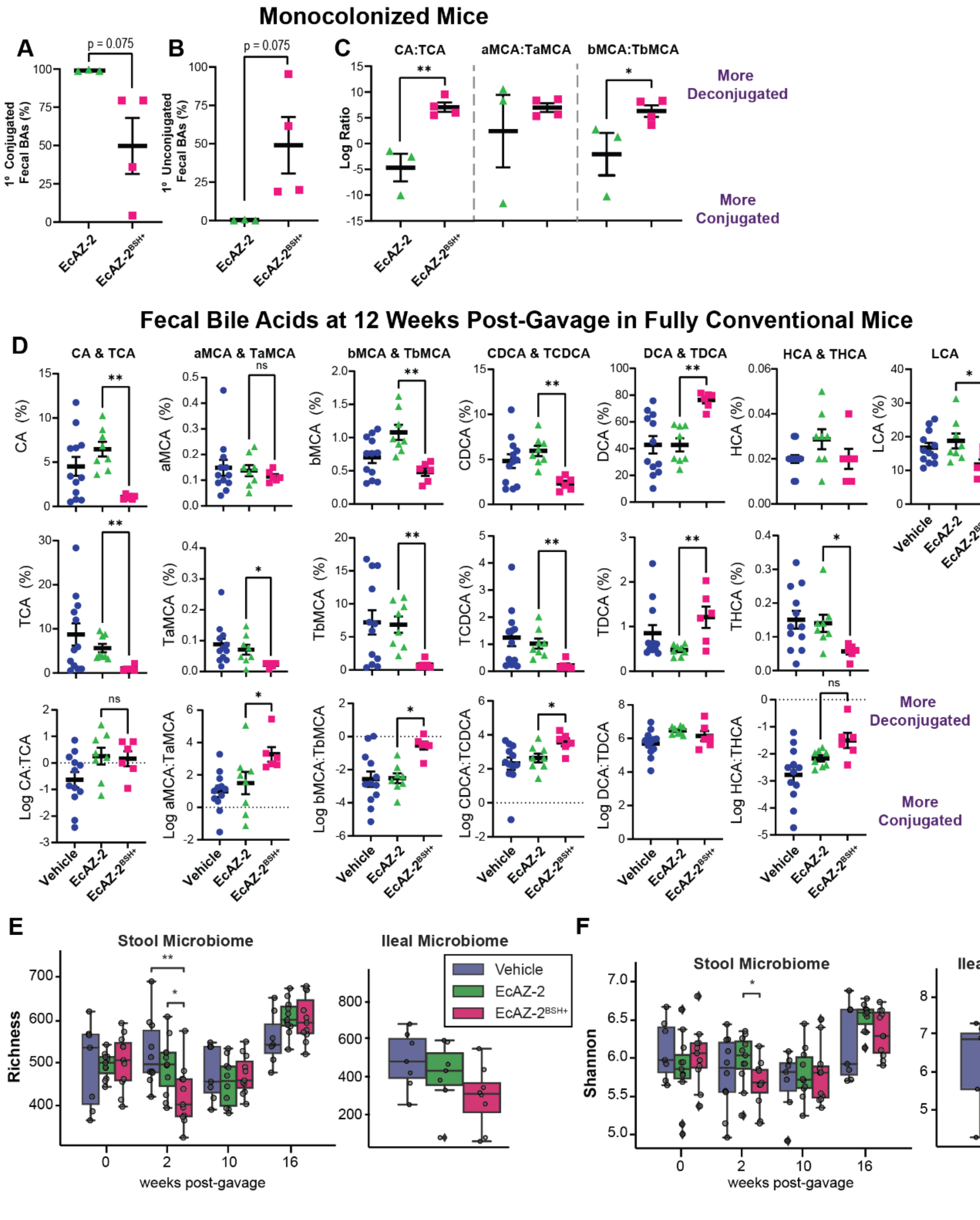


**Supplementary Figure 3 -**

**(A)** Primary conjugated fecal bile acids,
**(B)** Primary unconjugated fecal bile acids,

**(C)** $Log_2$ ratio of deconjugated to conjugated bile acids in gnotobiotic mice mono-colonized with either EcAZ-2 or EcAZ-2$^{BSH+}$. Significant differences determined by student's t-test with normality verified through Q-Q plot.

**(D)** Deconjugated (top), conjugated (middle) and the $log_2$ ratio of deconjugated to conjugated fecal bile acids in mice treated with vehicle, EcAZ-2 or EcAZ-2$^{BSH+}$. Significant differences determined by Kruskal-Wallis test with post-hoc Dunn's multiple comparison test comparing EcAZ-2 and EcAZ-2$^{BSH+}$.

**(E)** Richness, and

**(F)** Shannon index from 16S performed on stool samples collected pre-treatment and at 2-, 10-, and 16-weeks post-gavage (left) and from the terminal ileum samples at the time of euthanasia (right). Significant differences were determined by Kruskal-Wallis test with post-hoc Dunn's multiple comparison test comparing all three conditions.

### Supplemental Figure 4

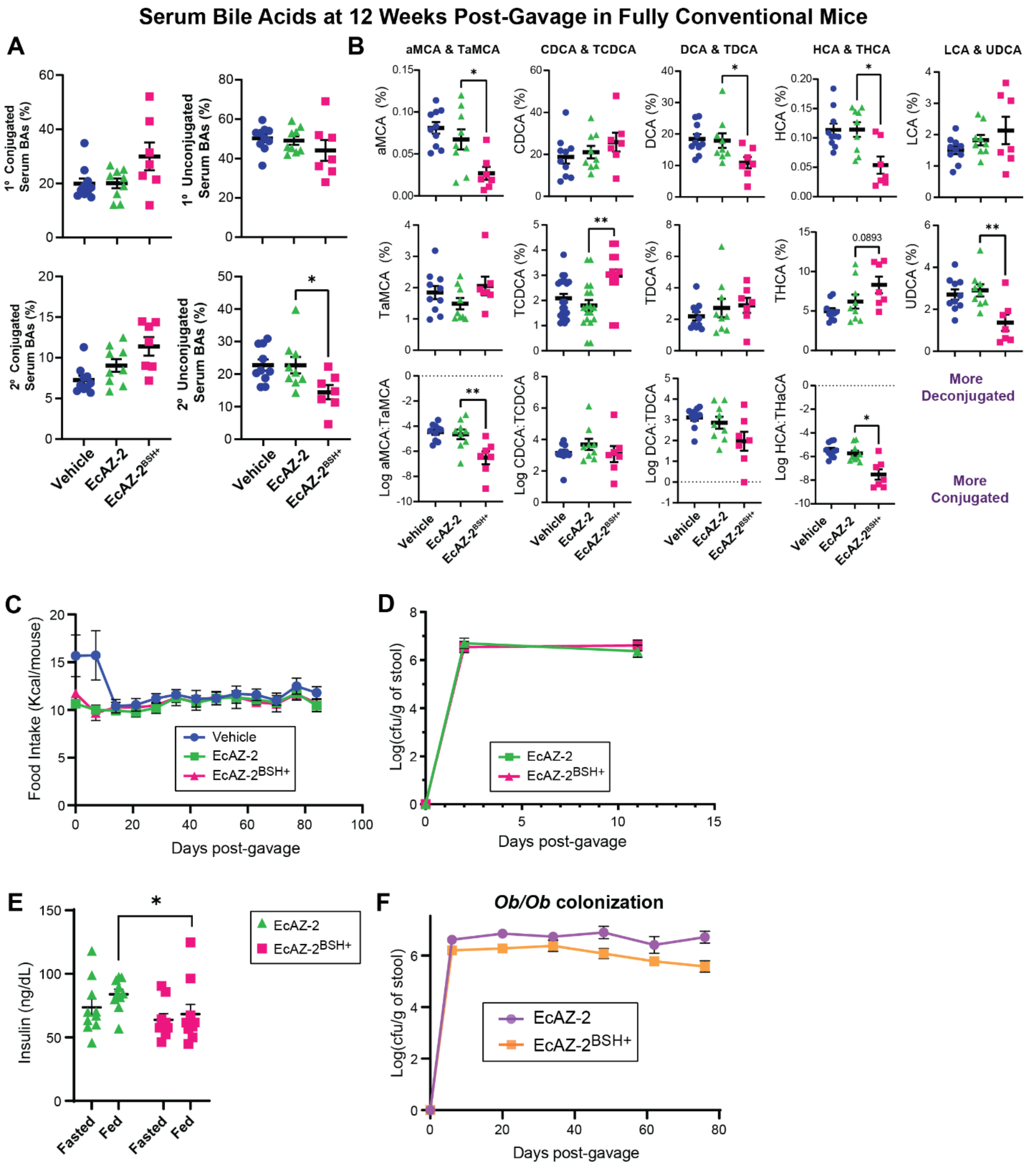


**Supplementary Figure 4 -**

**(A)** Primary conjugated, primary unconjugated, secondary conjugated, and secondary unconjugated serum bile acids from mice treated with a single gavage of vehicle, EcAZ-2 or EcAZ-$2^{BSH+}$ 12 weeks prior. Significant differences were determined by Kruskal-Wallis test with post-hoc Dunn's multiple comparison test comparing EcAZ-2 and EcAZ-$2^{BSH+}$.

**(B)** Deconjugated (top), conjugated (middle) and the $\log_2$ ratio of deconjugated to conjugated serum bile acids in mice treated with vehicle, EcAZ-2 or EcAZ-2$^{BSH+}$. Significant differences determined by Kruskal-Wallis test with post-hoc Dunn's multiple comparison test comparing EcAZ-2 and EcAZ-2$^{BSH+}$.

**(C)** Food intake in CR-WT mice treated with a single gavage of Vehicle, EcAZ-2, or EcAZ-2$^{BSH+}$.

**(D)** Colonization of EcAZ-2 and EcAZ-2$^{BSH+}$ after a single gavage in female CR-WT mice.

**(E)** Fasting (6 hours) and fed (30 min) insulin levels in female CR-WT mice treated with a single gavage of EcAZ-2 and EcAZ-2$^{BSH+}$ 22 days prior.

**(F)** Colonization of EcAZ-2 and EcAZ-2$^{BSH+}$ after a single gavage in *Ob/Ob* mice.
